# Drought degrades riparian subsidy quality and constrains aquatic ecosystem functioning

**DOI:** 10.64898/2026.07.10.737855

**Authors:** Rose M. Mohammadi, Albert Ruhí

## Abstract

The exchange of energy and organisms across habitat boundaries links aquatic and terrestrial ecosystems and sustains ecosystem functioning. Although disturbance may disrupt these linkages, the mechanisms at play remain poorly understood. Here, we investigated the extent to which flow intermittency may disrupt riparian-aquatic ecosystem linkages by altering consumer communities in the recipient ecosystem or by altering resource quality in the donor ecosystem. We ran an experiment in an intermittent river network in California, focusing on a critical forest-to-river subsidy (organic matter in the form of leaf litter), its transformation, and its reciprocal benefit (aquatic insect production). Using three riparian species (willow, cottonwood, and oak) at sites spanning a gradient of flow permanence, we quantified intraspecific plasticity in leaf traits (specific leaf area, nitrogen and phosphorus concentrations, and δ^13^C), measured decomposition rates, and estimated the secondary production of aquatic shredders (Plecoptera). Across all leaf species, decomposition rates were 16–36% lower at intermittent than perennial sites, an effect largely driven by intraspecific leaf trait plasticity rather than changes in consumer abundance. At high flow intermittency, willow experienced water stress (enriched δ^13^C) and reduced specific leaf area, while cottonwood showed primarily stoichiometric responses (reduced leaf nitrogen and phosphorus). Despite these divergent strategies, all species produced lower-quality litter at intermittent sites. Variance partitioning confirmed that initial litter quality uniquely explained 51.5% of variation in decomposition rates, more than double the contribution of invertebrate community metrics; and structural equation modeling revealed that both leaf traits and stonefly (Plecoptera) secondary production significantly predicted decomposition rates, with leaf traits exerting the stronger effect. Notably, stonefly secondary production was 37–98% lower at intermittent sites across leaf species. Because these insects later emerge as terrestrial adults, they provide a significant energy flux to riparian predators, and, thus, impoverished litter quality suppresses the reciprocal transfer of energy back to terrestrial food webs. As drought intensifies globally, the decoupling of terrestrial–aquatic linkages may begin in the riparian canopy.

## Introduction

Ecosystems are sustained by continuous flows of materials and organisms across habitat boundaries. These cross-ecosystem linkages, specifically the fluxes of organic and inorganic matter between terrestrial and aquatic systems, structure food webs and regulate landscape-scale ecosystem functioning (Nakano and Murakami 2001; Polis et al. 1997). Because recipient ecosystems often depend on externally derived resources from donor habitats (Polis and Hurd 1996; Wallace et al. 1997), changes in the quantity, timing, or quality of these subsidies can propagate far beyond their source. Disturbance and environmental stressors disrupt these linkages by altering resource transfers (Baxter et al. 2004). When subsidies arrive degraded or decoupled from consumer activity, their capacity to support subsequent processes may be fundamentally altered, with consequences that cascade through recipient ecosystems (Baxter et al. 2007; Fausch et al. 2010).

Aquatic ecosystems exemplify this connectivity. Rivers act as “biogeochemical reactors” that process terrestrial organic matter, a process central to the global carbon cycle (Battin et al. 2023). Leaf litter decomposition is an integrative ecosystem process that channels energy into aquatic food webs where it is incorporated into consumer biomass and thus secondary production (Wallace et al. 1997; Gessner et al. 1999). Decomposition rates depend on resource traits (e.g., leaf chemistry and morphology) (Pastor et al. 2014; García-Palacios et al. 2016), on the decomposer community (e.g., microorganisms and invertebrates) (Boyero et al. 2021; del Campo et al. 2025), and also on environmental conditions (e.g., water presence, temperature, pH) (Boyero et al. 2016). These drivers interact, and environmental stressors can simultaneously alter resource quality and decomposer activity, with complex joint effects on decomposition that make it difficult to predict ecosystem responses (García and Pardo 2015; Fenoy et al. 2020).

Global change is increasingly altering the quantity and quality of cross-ecosystem subsidies, rewiring the linkages between habitats. A growing body of work has examined how anthropogenic stressors operating within aquatic systems (e.g., pollutants, hydromorphological alteration, land use change, invasive species) modify subsidy dynamics and propagate effects into terrestrial food webs (Schulz et al. 2024; Twining et al. 2025). For instance, contaminants can reduce emergent insect biomass, limiting aquatic subsidies to riparian predators (Kraus et al. 2014), while altered flow regimes can disrupt the phenology and community composition of emergent insects (Leathers et al. 2024). However, a less explored pathway is how climatic water stress in the terrestrial donor ecosystem propagates through changes in subsidy quality before subsidies even enter the stream. When the donor ecosystem is itself stressed, it may deliver an impoverished resource that constrains functioning in the recipient ecosystem: a donor-mediated disruption that is mechanistically distinct from the aquatic-stressor pathways that have received the most attention to date.

River drying, which affects approximately 60% of river miles globally (Messager et al. 2021), disrupts litter decomposition through multiple pathways that have been well-documented. First, drying alters physical and microbial conditioning. As drying fragments the river continuum, isolated pools become hypoxic, accumulating toxic leachates that curb microbial activity even while leaves remain submerged (Dieter et al. 2011). When surface water disappears, physical abrasion and leaching slow, and solar radiation becomes the dominant driver, causing photodegradation and chemical alteration of the litter (del Campo et al. 2019). Second, drying acts as an environmental filter on decomposer communities. The absence of surface water reduces fungal and bacterial biofilm activity, preventing nutrient immobilization and rendering litter less palatable for invertebrates (Pastor et al. 2014). Drying fragments the river continuum, limiting the dispersal, abundance, and diversity of invertebrates even after flow resumption (Bogan et al. 2013; Datry et al. 2014; Sarremejane et al. 2020). Consequently, reduced decomposition rates in intermittent streams are commonly attributed to impoverished macroinvertebrate assemblages (Datry et al. 2011; del Campo et al. 2021).

However, this decomposer-centric perspective may overlook a concurrent pathway: alteration of the litter resource before it enters the stream. In dryland regions, such as Mediterranean-climate systems, riparian trees experience prolonged periods of reduced soil moisture (Quichimbo et al. 2020). During water stress, these trees can increase their intrinsic water use efficiency, the ratio of carbon acquired to water vapor losses via stomatal conductance, reflected in enriched leaf carbon isotope ratios (δ^13^C) (Stella and Battles 2010). Riparian trees also may exhibit morphological responses to water stress, including reduced leaf size and specific leaf area (SLA) (Stella and Battles 2010). Aridity can also reduce leaf nutrient concentrations (e.g., nitrogen and phosphorus) while increasing lignin and tannin content and leaf toughness (Rubio-Ríos et al. 2022). While such plasticity in leaf traits can strongly influence litter palatability and decomposability along gradients of water availability (LeRoy et al. 2007; Lecerf and Chauvet 2008; Boyero et al. 2017; Rubio-Ríos et al. 2022), its relative importance compared to drying-induced changes in decomposer communities remains unresolved in intermittent streams.

Here, we examine these two pathways, consumer-mediated *vs.* resource-quality-mediated, by which water stress alters leaf litter decomposition (Figure 1). We addressed four questions: (1) Does the decomposition of leaf litter differ in a predictable way in response to water stress across distinct riparian tree species? (2) Does water stress correspond to plasticity in leaf traits, resulting in altered litter quality? (3) Does water stress alter invertebrate community dynamics, specifically the secondary production of shredders? (4) Are decomposition rates more sensitive to hydrologically induced changes in leaf traits (the resource pathway) or changes in the consumer community (the consumer-driven pathway)?

**Figure 1.**
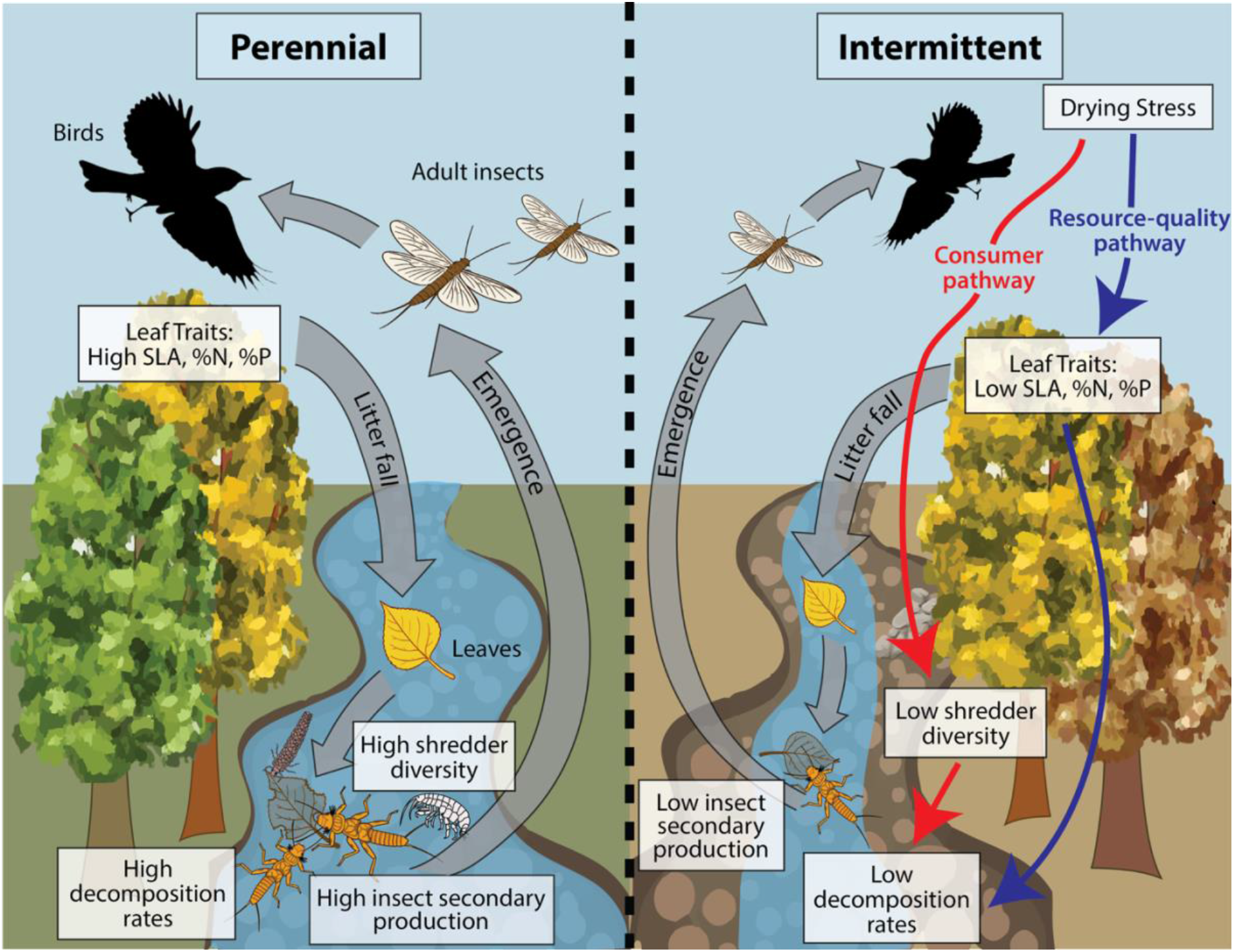
Conceptual model of how flow intermittency is expected to alter riparian–aquatic linkages through *consumer-mediated* and *resource-quality-mediated* pathways (red and blue arrows, respectively). Perennial reaches (left) are expected to support high quality leaf litter, high shredder diversity, and thus, high decomposition rates (*prediction 1*) and aquatic insect secondary production (*prediction 3*). In turn, in intermittent reaches (right), flow intermittency may alter riparian leaf functional traits (*prediction 2*; e.g., SLA, %N, %P), lowering litter quality (in blue, resource-quality-mediated pathway), and may reduce shredder diversity (in red, consumer-mediated pathway) and secondary production (*prediction 3*). While past work has largely focused on the *consumer-mediated* pathway, we predict the *resource quality-mediated* pathway may similarly influence decomposition rates (*prediction 4*).

We tested the general hypothesis that environmental stress controls organic matter breakdown by filtering resource quality and consumer community composition. We tested the following four predictions: (*Prediction 1*) Decomposition rates will increase with increasing flow permanence, with higher rates at more permanent reaches reflecting reduced water stress (Datry et al. 2011; del Campo et al. 2021). The magnitude of this increase would vary by tree species identity, with deciduous, non-sclerophyllous cottonwood and willow decomposing faster than evergreen, sclerophyllous coast live oak (Webster and Benfield 1986; LeRoy et al. 2007; Alonso-Forn et al. 2023). (*Prediction 2*) Water stress will induce intraspecific plasticity in leaf traits, with more permanent sites producing higher-quality litter (characterized by higher nitrogen and phosphorus concentrations and SLA) than intermittent sites (Stella and Battles 2010). This plasticity will vary among species, with cottonwood and willow exhibiting stronger increases in specific leaf area and nutrient concentrations in response to water stress than the oak (Chacon et al. 2020; Hultine et al. 2020). (*Prediction 3*) Shredder density and secondary production will increase with flow permanence (Datry et al. 2011; Bogan et al. 2013; Datry et al. 2014; Sarremejane et al. 2024), and litter bags containing higher-quality leaf species would support greater shredder colonization than those containing recalcitrant litter (Graça et al. 2015; Graça and Cressa 2010; Lecerf et al. 2007). (*Prediction 4*) Finally, we predicted that both litter quality (the resource pathway) and macroinvertebrate assemblages (the consumer pathway), each structured by water stress, would similarly explain variation in litter decomposition rates (García-Palacios et al. 2016).

## Methods

### Study site

Chalone Creek, Pinnacles National Park, California is an intermittent watershed where about ∼90% of the stream network naturally dries each year. Reaches span a range of flow permanence (∼15–100%). We used flow permanence as a proxy for water stress, classifying reaches as perennial (year-round surface flow; low stress) or intermittent (seasonally dry; high stress) for descriptive purposes. All statistical models and visualizations used flow permanence as a continuous variable (percentage of days with surface flow, ranging from 36–100% across study sites; Appendix S1: Table S1), to capture the full gradient of intermittency across sites. Our array of sites constitutes a quasi-experimental framework. Situated within a single small watershed, they differ in flow permanence while sharing climate, geomorphology, and riparian species composition, making hydroperiod the primary axis of variation among sites and limiting many confounds that would complicate broader cross-site comparisons (Bogan and Carlson 2018; Fournier et al. 2023). The watershed has a Mediterranean climate, with hot, dry summers and cool, moderately wet winters, and a long-term average annual precipitation of 414 mm (Western Regional Climate Center 2024a, 2024b). The riparian area is dominated by the deciduous *Salix laevigata* Bebb (red willow, “willow”), *Salix lasiolepis* Benth. (arroyo willow), *Populus fremontii* S. Watson (Fremont cottonwood, “cottonwood”), *Quercus lobata* Née (valley oak), and *Platanus racemosa* Nutt. (Western sycamore) along with the evergreen *Quercus agrifolia* Née (coast live oak, “oak”) *and Pinus sabiniana* Douglas (gray pine). The growing season typically ends in late October during non-drought years (Mohammadi et al. 2025).

### Decomposition experiment

In November 2023, we collected senesced leaves from the branches of riparian willow, cottonwood, and oak trees at 3 perennial and 4 intermittent reaches (Appendix S1: Figure S1). Five grams of air-dried leaves were enclosed in coarse (1 cm) and fine (250 μm) bags. Coarse mesh bags allowed invertebrates and microorganisms to enter and consume leaves (total decomposition), while fine mesh bags only permitted microbial decomposition (i.e., by fungi and bacteria). Paired coarse and fine mesh litter bags have known limitations: abrasion rates are higher in coarse mesh bags which can inflate apparent macroinvertebrate contributions to breakdown, and microbial respiration rates can differ between mesh sizes depending on leaf type and decomposition stage (Tomczyk et al. 2022). Fine mesh bag data were therefore not incorporated into the variance partitioning or structural equation models; they are presented descriptively to characterize decomposition dynamics. Leaf bags were deployed at their collection sites in January 2024 when both perennial and intermittent sites had surface water flow. Three replicate bags per species and mesh type were retrieved 28, 56, 84, and 112 days post-deployment. This local-origin deployment is realistic in our system, as riparian trees supply litter to their adjacent reach, and, as litter abscission at our sites peaks from October to November, the litter is incorporated into the channel during flow and flood pulses.

We evaluated six decomposition models for each site, species, and mesh size group (n=36) using the ‘litterfitter’ package (Cornwell and Weedon 2014) in R (R Core Team 2023). We selected the negative exponential model for all analyses to prioritize parsimony and comparability with existing literature (Olson 1963; Lecerf 2021). Details of decomposition model selection and alternative model fits are provided in Appendix S1 (Section S1, Figure S2).

To determine whether litter decomposition rates differed across the flow permanence gradient, and whether invertebrate access mediated these differences (Question 1), we modeled the natural log-transformed proportion of mass remaining as a function of *time* (days), *flow regime* (perennial vs. intermittent), and *mesh size* (coarse vs. fine). We fit a linear mixed-effects model using the ‘lme4’ and ‘lmerTest’ packages in R (Bates et al. 2015; Kuznetsova et al. 2017).

To investigate significant interaction terms involving *time* (which represent differences in decomposition rates, *k*), we performed Tukey-adjusted post-hoc pairwise comparisons of slopes using the ‘emmeans’ package (Lenth 2023). We compared decomposition rates among species averaged across other factors, and between flow regimes within each species to parse the significant three-way interaction.

### Leaf litter quality

We measured specific leaf area (SLA), the ratio of leaf surface area to mass (mm^2^/mg), for 30 senesced leaves per site-species combination using the LeafByte app to measure leaf area (Getman-Pickering et al. 2020) and an analytical balance. To estimate the initial chemical quality of senesced leaves, we homogenized 5 replicate samples per site and species, each consisting of 20 leaves. All leaves were oven-dried at 60°C and ground to a fine powder. Nitrogen (%N) and carbon (%C) concentrations were measured using a combustion system equipped with an induction furnace, thermal conductivity detector, and infrared detector. Phosphorus (%P) was quantified following a nitric acid/hydrogen peroxide microwave digestion using an Inductively Coupled Plasma Atomic Emission Spectrometry (ICP-AES). Stable carbon isotope ratio (δ^13^C) was determined for 3 of the 5 replicates using a PDZ Europa Scientific 20/20 mass spectrometer with a continuous-flow CN elemental analyzer. To test whether flow permanence was associated with intraspecific plasticity in leaf traits (Question 2), we evaluated differences in trait values across flow regimes using linear mixed-effects models for each species, with *flow permanence*, *leaf species*, and their interaction as fixed effects and *site* as a random effect. Species-specific slopes of trait responses to flow permanence were extracted and tested against zero.

### Invertebrate community

Invertebrates present in the coarse mesh bags were identified to family or genus level and categorized into functional feeding groups (Vieira et al. 2006). Stoneflies (Plecoptera) dominated the shredder community (62.5% abundance), with taxa particularly well adapted to Californian intermittent streams (Fournier et al. 2023, Bogan & Carlson 2018). We focused our secondary production efforts on the abundant family Capniidae, genus *Nemoura*, and the species *Malenka californica*, which were measured for body length. Biomass was then estimated by converting individual body lengths to dry mass using published length–mass relationships (Benke et al. 1999).

To test whether invertebrate community composition differed between flow regimes (Question 3), we used non-metric multidimensional scaling (NMDS) based on Bray-Curtis dissimilarities of Hellinger-transformed invertebrate abundance data. Ordinations were performed in three dimensions in the ‘vegan’ package (Oksanen et al. 2025). Differences in community composition between flow regimes were tested using permutational multivariate analysis of variance (PERMANOVA) with *flow regime* as a fixed factor and permutations (*n* = 999) constrained by sampling date to account for temporal non-independence.

To test whether flow permanence was associated with changes in shredder density and taxonomic richness (Question 3), we analyzed invertebrate count data using generalized linear mixed models (GLMMs) with a negative binomial distribution to account for overdispersion and zero inflation in the invertebrate count data. Models included *leaf species*, *flow permanence*, and their interaction as fixed effects, with *site* as a random intercept. *Flow permanence* was standardized (z-scored) prior to model fitting. Significance of fixed effects was determined using Type III Wald chi-square tests. GLMMs for collectors, scrapers, and predators are presented in Appendix S1: Figure S3 and Appendix S1: Table S3.

To test whether higher flow permanence was associated with an increase in Plecoptera secondary production (Question 3), we estimated benthic Plecoptera secondary production using the production to biomass (P/B) ratio method by multiplying seasonal biomass by a P/B ratio estimated from our data using the size-frequency method (Benke 1984). Secondary production estimates were aggregated to the site × species level, as the size-frequency method requires pooling individual body length measurements across all bags within a site, yielding a single production estimate per site × species combination (*n* = 17). To maintain statistical power, we grouped sites by flow regime type (perennial *vs.* intermittent). For each species, differences in production between flow regimes were assessed using 95% confidence intervals derived from 1,000 bootstrap iterations, which were used to characterize uncertainty in the production estimates. Two-tailed p-values were calculated from the proportion of bootstrap differences greater or less than zero.

### Drivers of decomposition

To disentangle the relative contributions of litter quality, invertebrate community structure, and environmental context to total decomposition rates (Question 4), we performed a variance partitioning analysis with three predictor matrices: (1) litter traits (initial SLA, %N, %P, %C); (2) invertebrate community metrics (densities of shredders, scrapers, predators, collectors, and parasites); and (3) environmental conditions (flow permanence [%], tributary, drainage area [m^2^], and average water temperature [°C] during the experiment; see Appendix S1: Section S2 for measurement details and Appendix S1: Figure S4 for time series). Adjusted R^2^ values were used to estimate the proportion of variance uniquely attributable to each predictor set and their shared fraction. We used partial redundancy analysis (RDA) to quantify the unique and shared variance in *k* explained by each matrix.

To evaluate the direction and relative magnitude of pathways by which flow permanence influences leaf litter decomposition (Question 4), we constructed a piecewise structural equation model (piecewise SEM) using the ‘piecewiseSEM’ package (Lefcheck 2016). Our models were designed to test two alternative but non-exclusive hypotheses: (1) a consumer-mediated pathway, whereby flow permanence reduces Plecoptera secondary production and thereby slows decomposition, and (2) a resource-mediated pathway, whereby drying alters leaf litter traits (e.g., nutrient concentrations and SLA) that subsequently constrain decomposition rates. Accordingly, the model included direct paths from flow permanence to Plecoptera secondary production and to litter quality traits, as well as paths from both Plecoptera secondary production and litter traits to decomposition rates. Leaf trait submodels were fit as linear models since flow permanence as a continuous site-level predictor absorbed between-site variance, rendering the random intercept redundant (confirmed by singular fits). Production and decomposition submodels were fit as linear mixed-effects models with site as a random intercept to account for residual clustering of species-level observations within sites. A correlated error term between %N and %P was included based on a significant directed separation test indicating residual covariance beyond their shared response to flow permanence. Directed separation tests indicated no significant residual covariance among litter traits after accounting for flow regime, suggesting that additional correlated error terms were not warranted. We assessed model fit using Fisher’s *C* statistic, and path coefficients standardized by standard deviation were used to compare the strength of competing pathways.

## Results

### Leaf litter decomposition dynamics

Decomposition rates (*k*) (Figure 2) increased significantly with flow permanence (Flow × Time, F_1,256_ = 6.15, *p* = 0.014) and differed among *species* (Species × Time, F_2, 257_ = 13.93, *p* = 0.021). Cottonwood and willow decomposed at similar rates (*p* = 0.950) and faster than the oak (*p* < 0.001 for both comparisons). The effect of flow permanence on decomposition rates varied by species (Flow × Species × Time, F_2,257_ = 5.09, *p* = 0.007): willow showed the strongest response, with decomposition rates ∼2.3× faster at perennial sites compared to the most intermittent site, while oak showed an intermediate response (1.4× faster). Cottonwood showed no significant relationship between decomposition rate and flow permanence (*p* = 0.60), and willow’s slope along the flow permanence gradient differed significantly from oak’s (*p* = 0.014). Although decomposition tended to be faster in coarse mesh bags than fine mesh bags, this difference was not significant (Mesh × Time, F_1, 256_ = 1.46, *p* = 0.228. No significant interaction was detected between *mesh size* and the other factors combined (Species × Flow × Mesh × Time, F_2,256_ = 0.763, *p* = 0.467), suggesting the responses to flow permanence were consistent across microbial-only and total decomposer communities.

**Figure 2.**
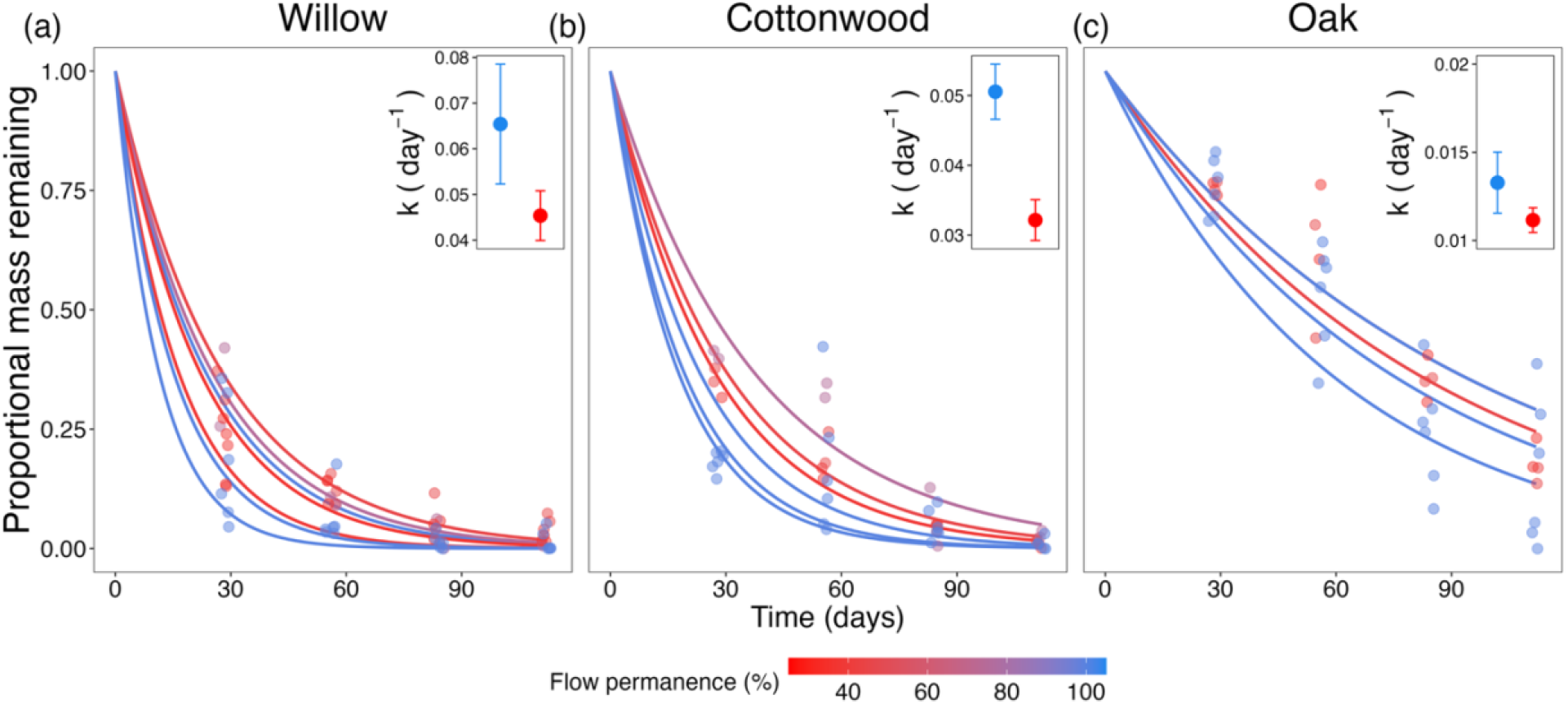
Litter decomposition dynamics for willow, cottonwood, and oak leaves. Sites with higher flow permanence, i.e. flowing year-round (blue) consistently showed faster decomposition rates than sites with lower flow permanence (red). Each line represents the fitted negative exponential decay model for a single site, colored by flow permanence at the site (%). Each point represents the raw data measurement (leaf mass remaining) from an individual coarse mesh bag jittered for visibility.

### The resource pathway: Environmental filtering of leaf litter quality

Leaf traits differed physiologically, morphologically, and chemically across flow permanence, though plasticity varied by species. Water use efficiency, measured via δ^13^C, declined significantly with increasing flow permanence across all three species (Figure 3a, F_1, 7.2_ = 11.69, *p* = 0.011), with no significant interaction between *flow permanence* and *species* (F_2, 50.5_ = 0.11, *p* = 0.893), indicating a consistent directional response to water stress regardless of species identity. Species-specific slope estimates were significant for willow (*p* = 0.015) and non-significant for cottonwood and oak (*p* = 0.061 and *p* = 0.062, respectively), though these slopes did not differ significantly from one another. Species differed in baseline δ^13^C values (F_2, 50.8_ = 4.55, *p* = 0.015). In contrast, there was no significant main effect of *flow permanence* on SLA (Figure 3b, F_1, 6.7_ = 2.75, *p* = 0.143), though there were species-level responses (Species × Flow permanence, F_2, 576_ = 38.24, *p* < 0.001). Willow leaves exhibited a significant positive relationship between flow permanence and SLA (*p* = 0.007), whereas cottonwood and oak showed weaker, non-significant SLA responses (*p* = 0.658, *p* = 0.727).

**Figure 3.**
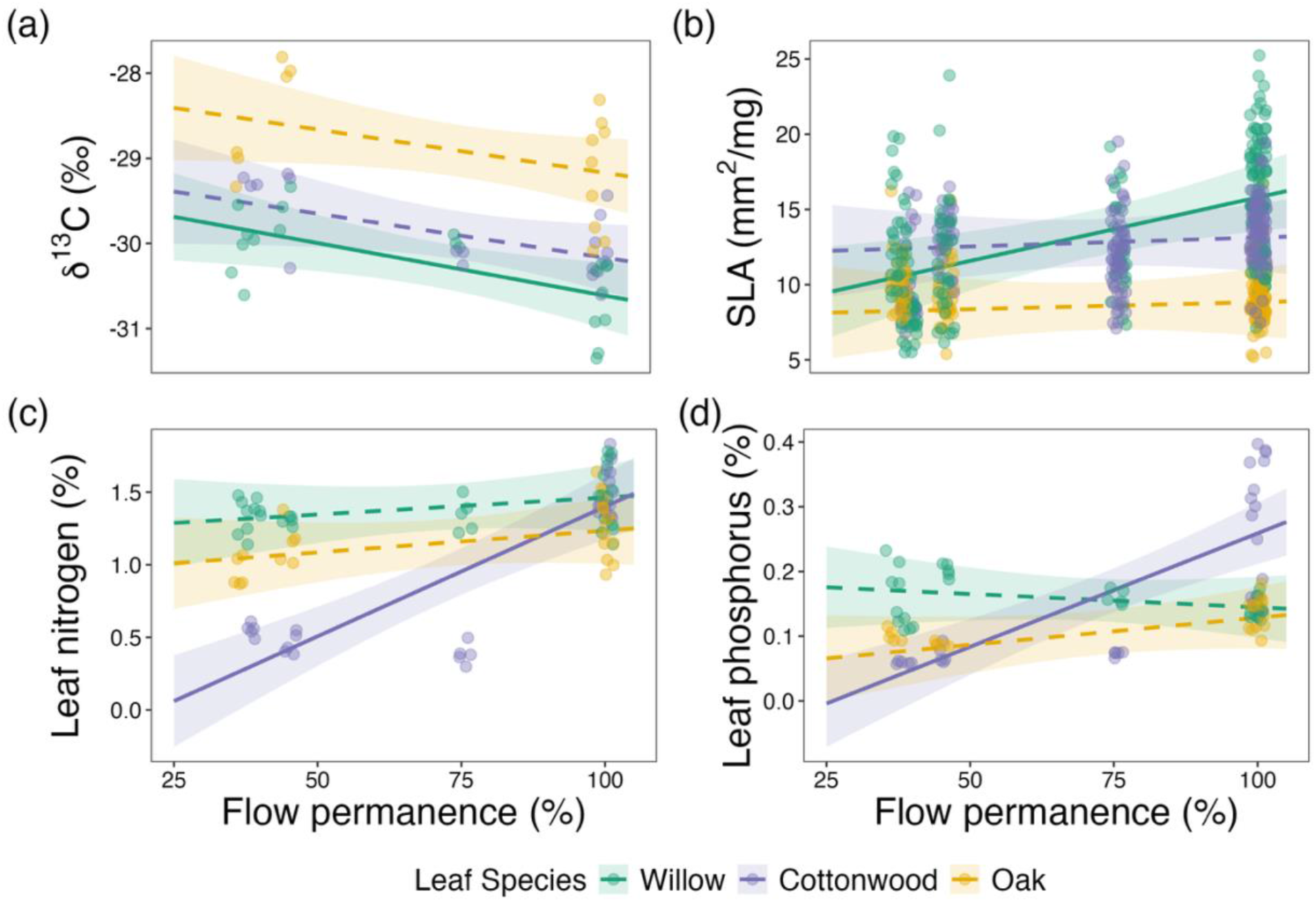
Relationships between flow permanence and riparian tree leaf traits. Leaves were collected from willow (green), cottonwood (purple), and oak (yellow) trees. Divergent species-specific strategies converge on low-quality litter at sites with reduced flow permanence. Panels display (a) stable carbon isotope ratios (δ^13^C, ‰), a proxy for water use efficiency; (b) specific leaf area (SLA, mm^2^/mg), a measure of leaf morphology and proxy for drought tolerance strategy; and (c-d) nitrogen content (%N) and phosphorus content (%P) respectively, reflecting litter nutritional quality. Lines represent predicted values from linear mixed-effects models and shaded ribbons indicate 95% confidence intervals. Solid lines indicate significant relationships between flow permanence and leaf traits within a species (*p* < 0.05, based on species-specific slope estimates; see Methods for details); dashed lines denote non-significant relationships. Points represent individual leaf measurements jittered for visibility.

Flow permanence and species interactively influenced leaf nutrient concentrations (Figure 3c, 3d), with significant main effects of both *flow permanence* (%N: F_1, 6.8_ = 13.76, *p* = 0.008; %P: F_1, 6.8_ = 9.33, *p* = 0.019) and *species* (%N: F_2, 85.6_ = 78.51, *p* < 0.001; %P: F_2, 86.2_ = 40.34, p < 0.001) as well as a significant flow permanence-by-species interaction (%N: F_2, 85.1_ = 51.05, *p* < 0.001; %P: F_2, 85.7_ = 49.05, p < 0.001). Cottonwood exhibited the strongest response to intermittency, with leaf %N and %P increasing significantly with flow permanence (both *p* < 0.001). Conversely, willow and oak showed no significant stoichiometric response to flow permanence (willow %N: *p* = 0.389, %P: *p* = 0.465; oak %N: *p* = 0.285, %P: *p* = 0.163).

### The consumer pathway: Invertebrate community composition and production

Invertebrate community composition differed significantly between perennial and intermittent reaches (Figure 4a; PERMANOVA, F_1, 136_ = 2.15, R^2^ = 0.016, p = 0.009). Shredder density increased significantly with flow permanence (Appendix S1: Figure S3; *χ*^2^(1) = 5.00, *p* = 0.025). The relationship did not differ among leaf species (*χ*^2^(2) = 0.245, *p* = 0.885). Shredder taxonomic richness showed a positive but non-significant trend with flow permanence (Appendix S1: Figure S3; *χ*^2^(1) = 2.39, *p* = 0.122). Richness did differ significantly among leaf species (*χ*^2^(2) = 7.76, *p* = 0.021), with cottonwood bags supporting significantly higher richness (2.16 taxa per bag) than willow bags (1.27 taxa per bag; *p* = 0.018), while oak bags (1.47 taxa per bag) did not differ significantly from willow or cottonwood (*p* = 0.786; *p* = 0.192).

**Figure 4.**
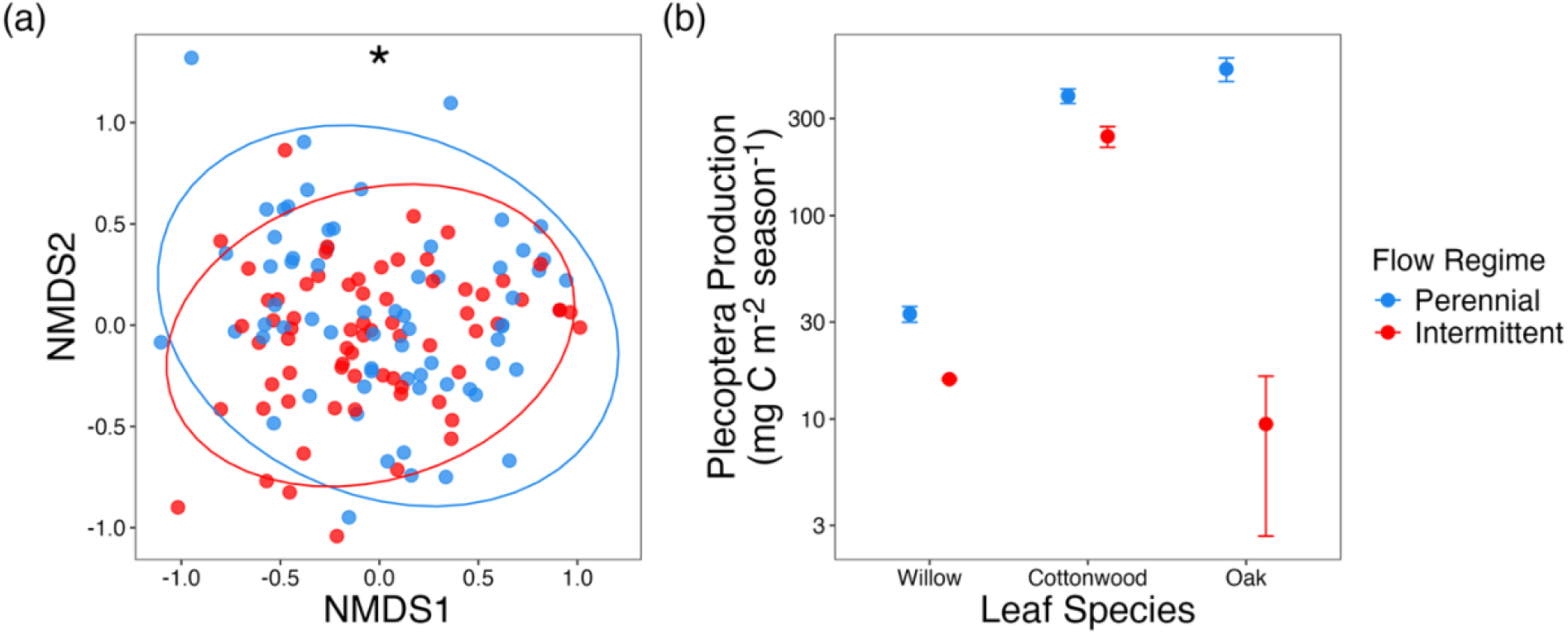
Flow permanence structures invertebrate communities and constrains Plecoptera secondary production. (a) Non-metric multidimensional scaling (NMDS) ordination of macroinvertebrate community composition, with points colored by flow regime (perennial in blue and intermittent in red) and ellipses indicating group dispersion. Community composition differed significantly between flow regimes (asterisk denotes significant grouping). (b) Mean Plecoptera secondary production (mg C m^−2^ season^−1^, log10 scale) at perennial (blue) and intermittent (red) sites by leaf species. Data are integrated across the entire decomposition period. Points represent bootstrap-estimated means (*n* = 1,000 iterations) and error bars represent the 95% confidence intervals of the bootstrap distribution.

Flow permanence exerted a strong constraint on Plecoptera secondary production (Figure 4b). Production was significantly lower at intermittent sites for all three leaf species (*p* < 0.001 for all pairwise comparisons). Intermittency reduced production by 37% for cottonwood, 52% for willow, and 98% for oak relative to perennial sites.

### Influence of leaf litter quality and shredder community on decomposition

Variance partitioning analysis revealed initial litter quality (SLA, %N, %P, and %C) was the dominant driver of total leaf litter decomposition (coarse *k*), uniquely explaining 51.5% of the explained variation in decomposition rates and accounting for a significant portion of variation when tested independently (F_4,13_ = 5.83, *p* = 0.008; Figure S5). The invertebrate community (functional feeding group densities) and environmental variables (flow permanence, tributary, water temperature, and drainage area) explained smaller unique fractions of variation (15.2% and 13.0%, respectively) and did not explain significant additional variation when considered independently (invertebrates: F_5,12_ = 1.14, *p* = 0.372; environment: F_4,13_ = 0.38, *p* = 0.826). There was a substantial shared component between leaf quality and invertebrates (13.9%), suggesting that high-quality litter resources supported higher shredder densities, collectively accelerating decomposition. Additionally, there was a smaller shared variance between environmental conditions and leaf quality (9.1%), likely reflecting that perennial flow regimes supported higher quality litter (e.g., higher %N). Collectively, all predictor sets combined explained 77.6% of total variation in decomposition rates (adjusted R^2^ = 0.776).

Piecewise SEM revealed that flow permanence exerted a strong bottom-up control on decomposition through the resource pathway (Figure 5). Higher flow permanence significantly predicted greater %N (standardized β = 0.502, *p* = 0.034), with additional positive but non-significant effects on %P (standardized β = 0.402, *p* = 0.098) and SLA (standardized β = 0.36, *p* = 0.141). Flow permanence also significantly predicted Plecoptera secondary production (standardized β = 0.587, *p* = 0.043), indicating that sites with higher flow permanence supported greater invertebrate production independent of litter quality. These traits, in turn, were strong predictors of decomposition rates: both SLA (standardized β = 0.966, *p* < 0.001) and %N (standardized β = 0.525, *p* < 0.001) had significant positive effects on total *k*. Secondary production was also significantly positively associated with decomposition rates (standardized β = 0.327, *p* = 0.004), consistent with a consumer-mediated pathway operating alongside the resource quality pathway. Leaf nitrogen and phosphorus showed significant residual covariance beyond their shared response to flow permanence (*r* = 0.514, *p* = 0.017). The model fit the data well based on the Fisher’s *C* statistic (C_4_ = 3.02, *p* = 0.554), indicating no significant missing pathways; and overall, the model explained 96% of the variation in decomposition rates (conditional R^2^ = 0.96).

**Figure 5.**
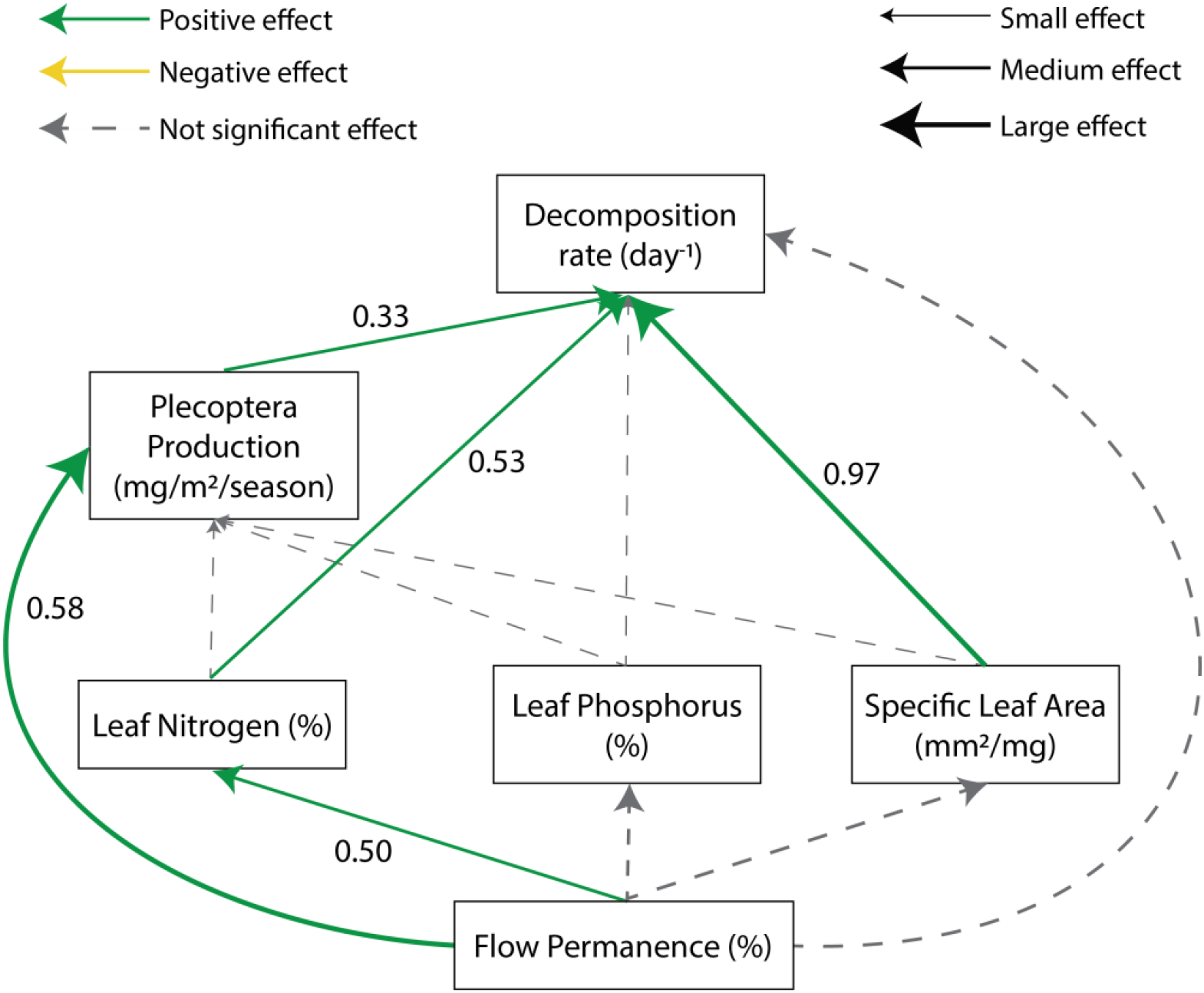
Flow permanence exerts control on leaf litter decomposition through litter quality and secondary production. Arrows represent hypothesized causal paths; solid lines indicate significant relationships (*p* < 0.05) and gray dashed lines indicate non-significant pathways included in the model. Arrow width is proportional to the standardized regression coefficient (shown numerically). Green arrows indicate positive relationships and yellow arrows negative effects.

## Discussion

Although ecosystems are linked by reciprocal exchanges of energy and materials, our understanding of how environmental stress disrupts these linkages and how such disruption propagates across coupled ecosystems remains incomplete. River networks span environmental gradients and are tightly linked with their watersheds, making them an ideal model system. Here, we showed that contrary to previous findings, flow intermittency can inhibit stream functioning not primarily by changing decomposer composition and abundance, but by imposing water stress on the riparian canopy, degrading the quality of the terrestrial litter subsidy at its source. Across three riparian tree species, intermittency consistently reduced decomposition rates, an effect driven largely by water stress in the riparian zone that produced nutrient-poor, recalcitrant litter. While shredders played a role, evidenced by the significant effect of Plecoptera secondary production on decomposition rates, variance partitioning and structural equation modeling identified initial litter quality as the dominant control on breakdown rates. These findings highlight a cross-ecosystem legacy effect: stress in a subsidizing ecosystem (the riparian forest) constrains ecosystem functioning in the recipient ecosystem (the stream), ultimately diminishing secondary production and the reciprocal transfer of energy back to terrestrial consumers. Our results align with growing evidence that global change is rewiring ecosystems not only by altering species interactions within habitats, but also the cross-ecosystem transfers that link them (Leathers et al. 2024; Ward et al. 2025).

### Resource quality supersedes shredder dynamics

A central finding of this study is that the reduction in decomposition rates in intermittent reaches was driven more by the physiological response of riparian trees to water stress than by shredder invertebrates. Though previous work has largely emphasized how drying events kill or displace aquatic invertebrates (a top-down constraint) (Datry et al. 2011, 2014; del Campo et al. 2021; Sarremejane et al. 2024), our variance partitioning analysis revealed that initial litter quality uniquely explained nearly half of the variation in decomposition rates; more than double the explanatory power of invertebrate assemblage metrics. The SEM further confirmed this bottom-up mechanism, with flow permanence strongly predicting leaf nitrogen and Plecoptera secondary production, and leaf traits exerting the strongest direct effects on decomposition. The litter quality pathway was stronger than the consumer pathway even after accounting for the significant positive effect of secondary production on decomposition rates.

Water stress altered leaf traits through species-specific strategies that converged on a common functional outcome: the production of low-quality litter. Willow responded to intermittency through morphological and physiological plasticity, producing leaves with significantly lower SLA and higher water-use efficiency (δ^13^C enrichment) as flow permanence declined. This reduction in SLA is consistent with a shift toward more xeromorphic leaf strategies and aligns with the leaf economics spectrum, in which lower SLA tends to covary with thicker, denser leaves and lower maximum leaf conductance in drought-tolerant species (Poorter et al. 2009; Wright et al. 2004). Concurrently, increasing water use efficiency is a common response of water-stressed plants as they close their stomata in response to higher vapor pressure deficits (Ehleringer and Osmond 2000), and has been observed in riparian Salicaceae seedlings under experimental (Stella and Battles 2010) and natural settings (Leffler and Evans 1999).

Cottonwood exhibited a stoichiometric response to flow intermittency. While previous studies have found that *Populus* also exhibited morphological trait plasticity (lower SLA) under water stress (Busch and Smith 1995; Stella and Battles 2010), our trees maintained similar morphology across flow regimes but significantly reduced leaf nitrogen and phosphorus concentrations. The decrease in leaf nutrient concentrations was likely driven by lower soil moisture which can limit nutrient supply through reduced mineralization and limited nutrient diffusion and mass flow in the soil (He and Dijkstra 2014). Both cottonwood and oak showed modest, non-significant δ^13^C enrichment, suggesting a milder water stress response to that of willow, though other studies of cottonwoods and Mediterranean evergreen oaks have documented shifts in δ^13^C during periods of drought (Stella and Battles 2010; Vaz et al. 2010). This divergence likely stems from interspecific differences in rooting depth; cottonwood and oak are reported to attain deeper maximum rooting depths (2.8 m and 10.7 m, respectively) than willow (1.5 m), which may increase access to deeper groundwater during the dry season (Rohde et al. 2024).

Despite these divergent strategies, the outcome was consistent: low-quality litter inputs that render detritus more recalcitrant for both microbial decomposers (Ferreira et al. 2006; Tuchman et al. 2002) and shredders (Graça and Cressa 2010; Lecerf et al. 2007), limiting their assimilation, activity, and growth (Graça et al. 2015). Our SEM results underscore the dominance of this resource constraint as leaf traits were strong direct predictors of decomposition rates independent of shredder secondary production, suggesting that reduced litter quality primarily slows breakdown by inhibiting microbial conditioning or physical fragmentation. Our findings mirror patterns documented across ecosystems, where drought slows most ecosystem processes (Knapp et al. 2024), including decomposition in grasslands and forests, primarily by altering producer traits and substrate quality rather than by directly removing consumers (Pugnaire et al. 2019; Seres et al. 2022).

### Flow intermittency as a key variable

Flow regime is widely recognized as a dominant driver governing instream community structure and function (Poff et al. 1997) as well as the important connections between river biodiversity and ecosystem processes (Palmer and Ruhi 2019). Our findings support this view and emphasize its relevance beyond the stream channel, showing that flow intermittency simultaneously structures processes across the riparian–stream interface. Riparian trees, which are spatially fixed and are dependent on shallow groundwater (Rohde et al. 2021), integrate water stress over long timescales. Once shed, their leaves are transported downstream, effectively propagating the legacy of water stress through the river network. These dynamics imply temporal lags between water stress in the riparian canopy, litter abscission, and subsequent downstream processing such that water stress can influence stream functioning well after the period of drying.

Although riparian vegetation is frequently included as a response variable in flow–ecology studies (Lytle and Poff 2004), it is rarely treated as a dynamic driver of instream functioning whose trait variation can influence downstream processes. While previous studies have examined how interspecific variation in leaf traits (Lecerf et al. 2007; Lecerf and Chauvet 2008; Leroy and Marks 2006) and drying-influenced preconditioning of litter (del Campo et al. 2019; Dieter et al. 2011) shape litter quality, our results indicate that flow intermittency drives significant intraspecific variation by morphologically and chemically altering the living riparian canopy before leaf fall. These results highlight the need to consider impacts of drought and low flows beyond the aquatic habitat, accounting for riparian trait responses that can propagate downstream and shape instream ecosystem functioning.

### Disrupted cross-ecosystem linkages and reciprocal subsidies

The consequences of altered leaf litter quality extend beyond the stream channel, with the potential to disrupt reciprocal subsidies back to terrestrial ecosystems. Both shredder densities and Plecoptera secondary production decreased with increasing flow intermittency, indicating an energetic constraint imposed by reduced resource quality. Aquatic insects that emergence as terrestrial adults represents a major energy pathway from streams to riparian food webs, providing prey for birds (Murakami and Nakano 2002), bats (Power and Rainey 2000), lizards (Sabo and Power 2002), and spiders (Kato et al. 2003) that contributes 25–100% of the energy or carbon to such terrestrial consumers (Baxter et al. 2005). Adult emergence biomass represents, on average, ∼19% of aquatic insect secondary production (Gratton and Zanden 2009). Thus, reduced secondary production implies a diminished return flux of energy to the terrestrial food webs.

Disrupted feedback loops may represent an underappreciated consequence of global change. Donor-mediated disruptions to cross ecosystem linkages have been observed in other systems, including in coastal sandy-beach ecosystems where declines in kelp and seagrass reduce wrack subsidies that fuel microbial processing and nutrient regeneration, thereby weakening the return flux of dissolved nutrients from beaches to coastal waters (Hyndes et al. 2022); agro-ecosystems where deer transfer nutrients from fertilized croplands to forest patches, altering forest nutrient availability in ways that can feedback to plant nutrient content and palatability and the quality of the resources subsequently available to herbivores (Abbas et al. 2012); and streams where reduced terrestrial invertebrate inputs triggered a trophic cascade that suppressed the reciprocal flux of adult insects back to the riparian forest (Baxter et al. 2004). Our study adds to this growing evidence by showing that environmental stress can impoverish cross-ecosystem subsidies through changes to the subsidy quality alone. As flow intermittency increases in prevalence due to climate change (Ayers et al. 2024; Carlson et al. 2024), such trait-mediated disruptions of reciprocal subsidies may decouple habitats that have historically functioned as integrated meta-ecosystems.

### Limitations and future directions

While our results strongly support a litter quality-mediated control over decomposition rates, several mechanisms warrant further investigation. First, although our analyses indicate that decomposition rates were more strongly constrained by litter quality than by consumer production, consistent with microbially mediated control of litter breakdown, we did not directly quantify microbial biomass or activity (e.g., via ergosterol concentration). Future work measuring microbial colonization and function on stress-altered litter would clarify how drying impacts fungal- and bacterial-driven decomposition. Second, by deploying leaf litter locally, our experiment isolated donor effects but did not capture the downstream transport and mixing of litter that occurs naturally in river networks (Catalán et al. 2023). Flow intermittency creates a habitat mosaic, allowing litter produced under different flow regimes to mix within individual reaches. Tracking litter transport across the stream network would help resolve how hydrologic connectivity modulates the spatial distribution of altered subsidies.

Notably, this study took place during an above-average precipitation year (∼40% above the long-term average), suggesting that even seasonal drying can substantially alter riparian leaf traits and alter cross-ecosystem subsidies. Under projected increases in drought frequency, duration, and intensity in many regions of the world (Padrón et al. 2020), such trait-mediated effects may intensify or compound over time, amplifying their effect on ecosystem functioning. Future work should evaluate how the strength and persistence of these responses scale with drought severity.

Our study demonstrates that stream drying disrupts stream–riparian linkages through a donor-mediated pathway that originates on land. By imprinting water stress on riparian vegetation, flow intermittency delivers an impoverished litter subsidy to the stream that constrains processing and suppresses the secondary production of aquatic consumers, likely diminishing the reciprocal flow of energy back to terrestrial predators. These results underscore the importance of a meta-ecosystem perspective, which recognizes that ecosystem functioning depends on interactions among coupled habitats (Bond and Chase 2002; Loreau et al. 2003).

Considering habitats in isolation risks overlooking mechanisms through which stress propagates across boundaries (Larsen et al. 2016), particularly when donor responses alter subsidy quality (Osakpolor et al. 2023). Overall, our findings reveal that subtle changes in donor ecosystems can generate disproportionate, cross-boundary consequences, exposing a key vulnerability of coupled ecosystems to global change.

## Acknowledgements

This research was supported by a National Science Foundation (NSF) CAREER award to A.R. (DEB-2047324) and by the Miller Institute for Basic Research in Science, UC Berkeley. R.M.M. was funded by the NSF Graduate Research Fellowship Program. This work was conducted on the ancestral lands of the Amah Mutsun and Chalon peoples. We thank Wildlife Biologist Paul G. Johnson and the Pinnacles National Park staff. We also thank Kendall Archie and the undergraduate researchers from the Undergraduate Research Apprentice Program for their help in the field and lab.

## Author contributions

R.M.M. and A.R. conceptualized the study. R.M.M. collected and analyzed the data and wrote the first draft of the manuscript. A.R. acquired funding and reviewed and edited the manuscript.

## Conflict of Interest Statement

The authors declare no conflicts of interest.

## Notes

### Competing Interest Statement

The authors have declared no competing interest.

