## Appendix S1 for "Drought degrades riparian subsidy quality and constrains aquatic ecosystem functioning"

### Drought degrades riparian subsidy quality and constrains aquatic ecosystem functioning

**Rose M. Mohammadi[1]* and Albert Ruhí[1]**

#### Section S1: Decomposition model selection

We compared the negative exponential, negative exponential model with a non-zero asymptote, Weibull, discrete parallel, discrete series, and continuous quality models. We used the package ‘litterfitter’ (Cornwell and Weedon 2014) in R (R Core Team 2023) to fit each time series to the analytical models. ‘Litterfitter’ finds parameters that minimize the negative log-likelihood for a given dataset and model. We manually adjusted the parameter space for each model optimization until all optimized parameter values were off the boundaries, with a minimum parameter value of $1 \times 10^{-20}$ (Robbins et al. 2025). We ran 9999 iterations for each model to each time series and checked convergence. We compared model fits using Akaike’s Information Criterion corrected for small sample size (AICc).

The Weibull model was the best-fitting or effectively equivalent model ($\Delta AICc \leq 2$) in 29 of 36 groups, while the negative exponential model also provided an equivalent fit in the majority of cases (25 of 36 groups). Weibull half life and mean residence time estimates are shown in Figure S2.

The negative exponential model follows the form:

$$M_t = M_0 e^{-kt}$$

where $M_t$ is the remaining mass at time $t$, $M_0$ is the initial mass, $k$ is the decomposition rate, and $t$ is the time in days.

#### Section S2: Stream temperature measurements

Two temperature sensors were deployed at the upstream and downstream ends of each study reach. Five sites had overlapping records from both sensors, while two sites had data from only a single sensor throughout the deployment period. For sites with overlapping records, upstream and downstream temperatures were highly concordant (mean Pearson's $r = 0.96$). Mean temperature differences between sensors were near zero (average of 0.30°C). Accordingly, temperature records were pooled to characterize reach-scale conditions, and data from the remaining sensor were used when only one sensor was available.

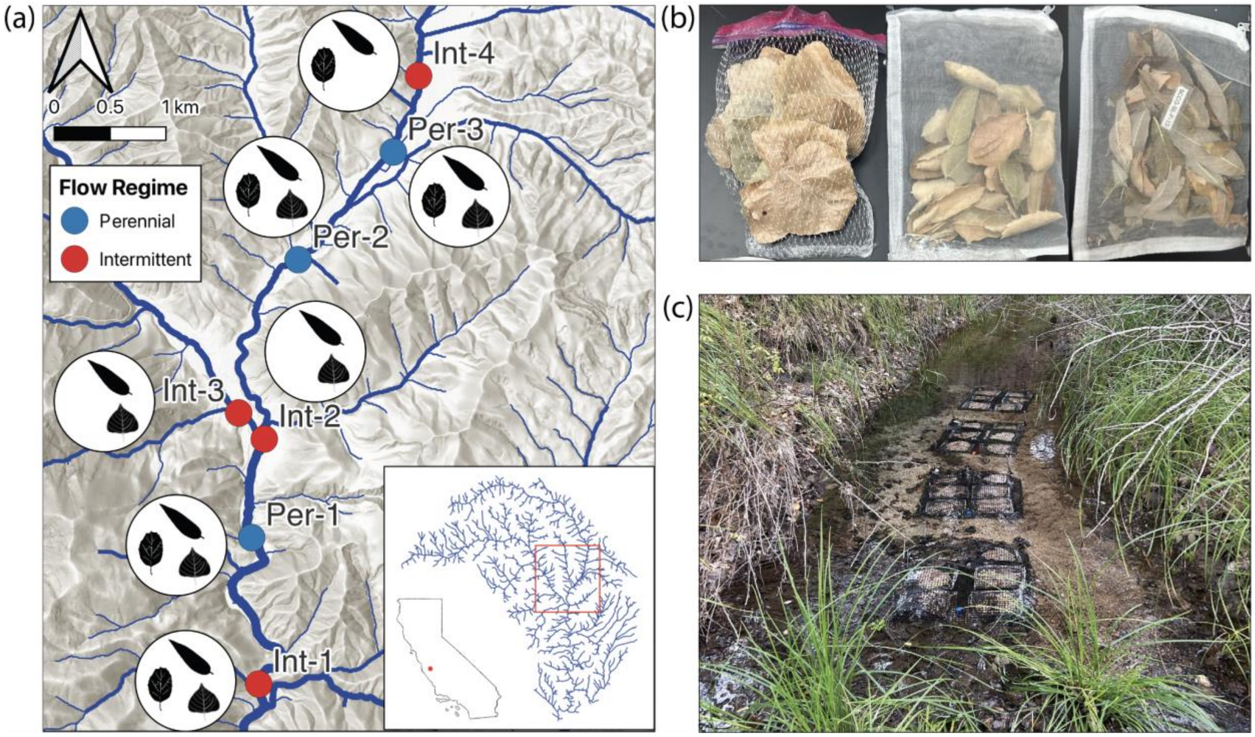


**Figure S1. Map of the 7 study sites in Chalone Creek, Pinnacles National Park, CA.** (a) Sites were categorized as perennial ("Per," blue, *n*=3) or intermittent ("Int," red, *n*=4) based on continuous flow monitoring since 2015. White circles next to the sampling sites contain leaf silhouettes that denote the tree species from which leaf litter was collected at that location: willow (top), cottonwood (right), and oak (left). The inset map (bottom right) shows the location of the sites within the watershed as well as the location of the study region within California, USA. (b) Examples of litter bags used in the experiment, showing coarse-mesh (left, cottonwood leaves), which allowed macroinvertebrate access and fine-mesh (middle, oak leaves, and right, willow leaves), which excluded macroinvertebrates. (c) A representative deployment of litter bags (at Per-3) secured to the streambed.

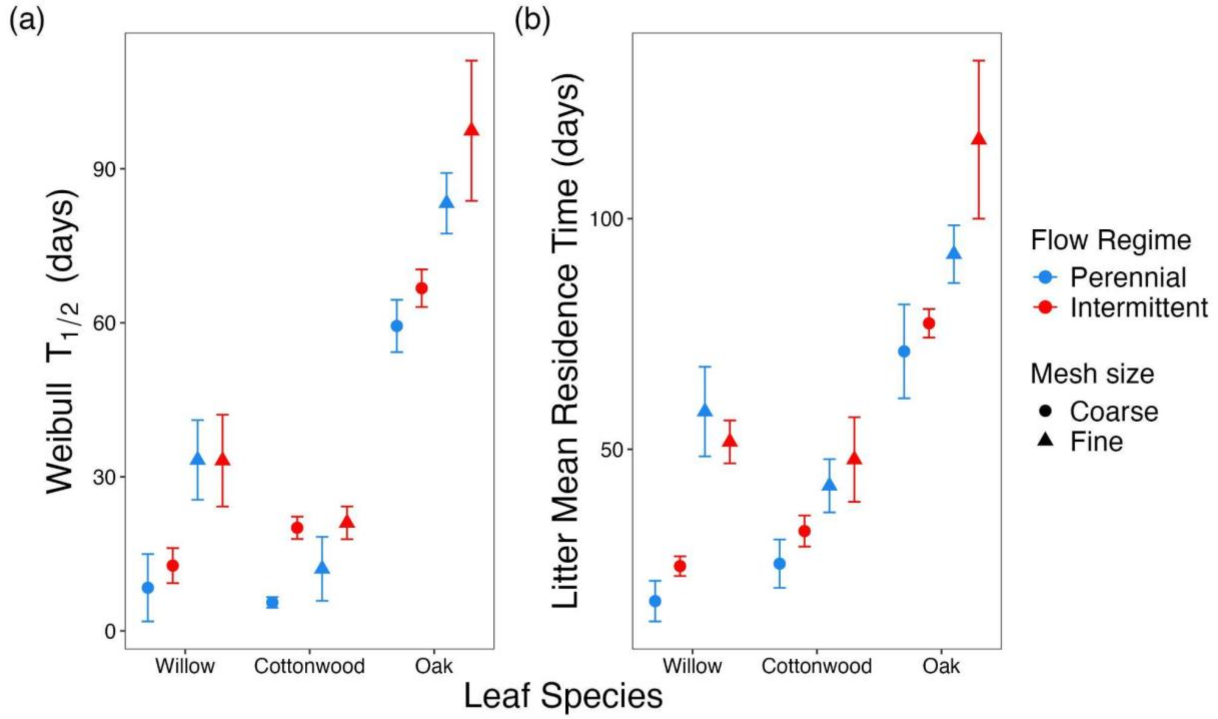


**Figure S2. Comparison of estimated leaf litter decomposition metrics derived from Weibull model fits.** Panels display (a) the Weibull half-life ($T_{1/2}$, days) representing the time required for 50% mass loss, and (b) the litter mean residence time (days) for willow, cottonwood, and oak. Points represent means grouped by perennial (blue) and intermittent (red) flow regimes and coarse (circles) and fine (triangles) mesh sizes. Error bars indicate standard error (SE).

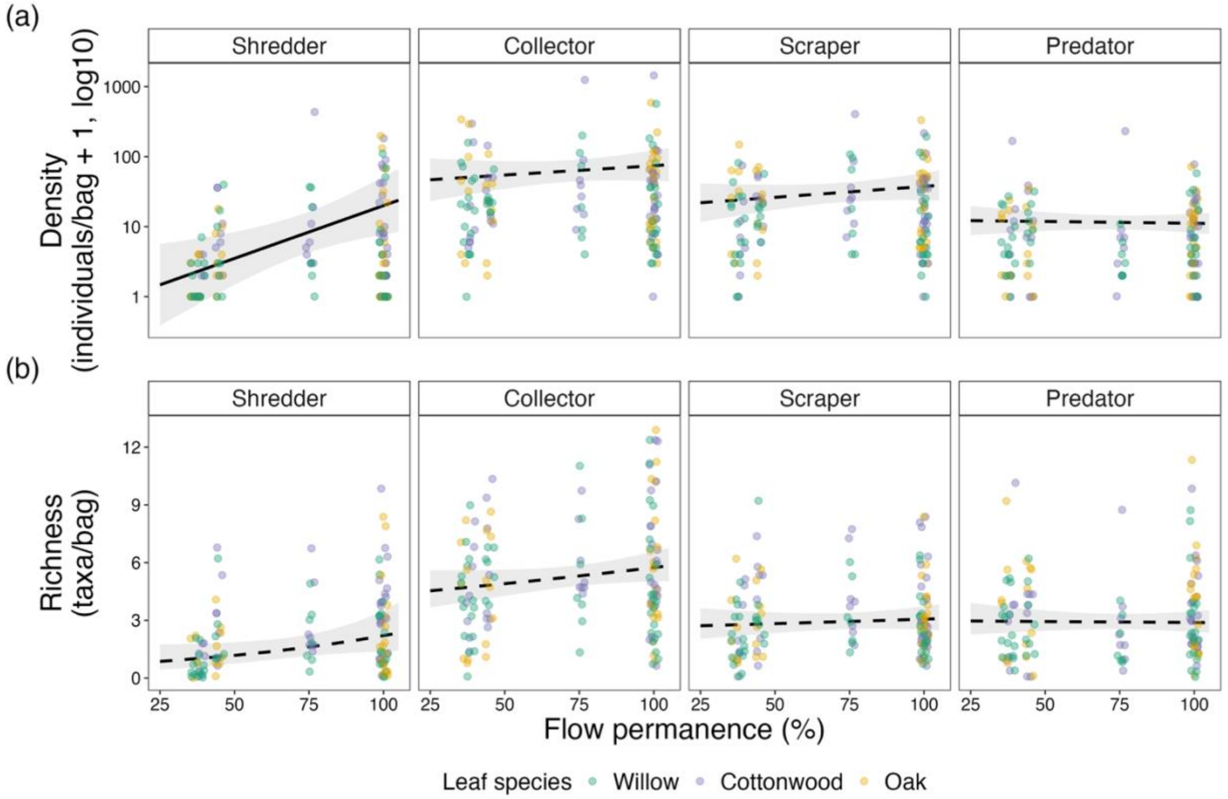


**Figure S3.** Density (individuals per coarse mesh bag + 1, log10 scale; top row) and taxonomic richness (taxa per bag; bottom row) of shredders, collectors, scrapers, and predators associated with willow, cottonwood, and oak leaf litter across a gradient of flow permanence. Lines represent predicted values from negative binomial generalized linear mixed-effects models (GLMMs) averaged over leaf species, with shaded ribbons indicating 95% confidence intervals. Solid lines indicate significant relationships between flow permanence and leaf traits within a species ($p < 0.05$, based on species-specific slope estimates); dashed lines denote non-significant relationships (Table S3). Points represent individual bag-level observations colored by leaf species.

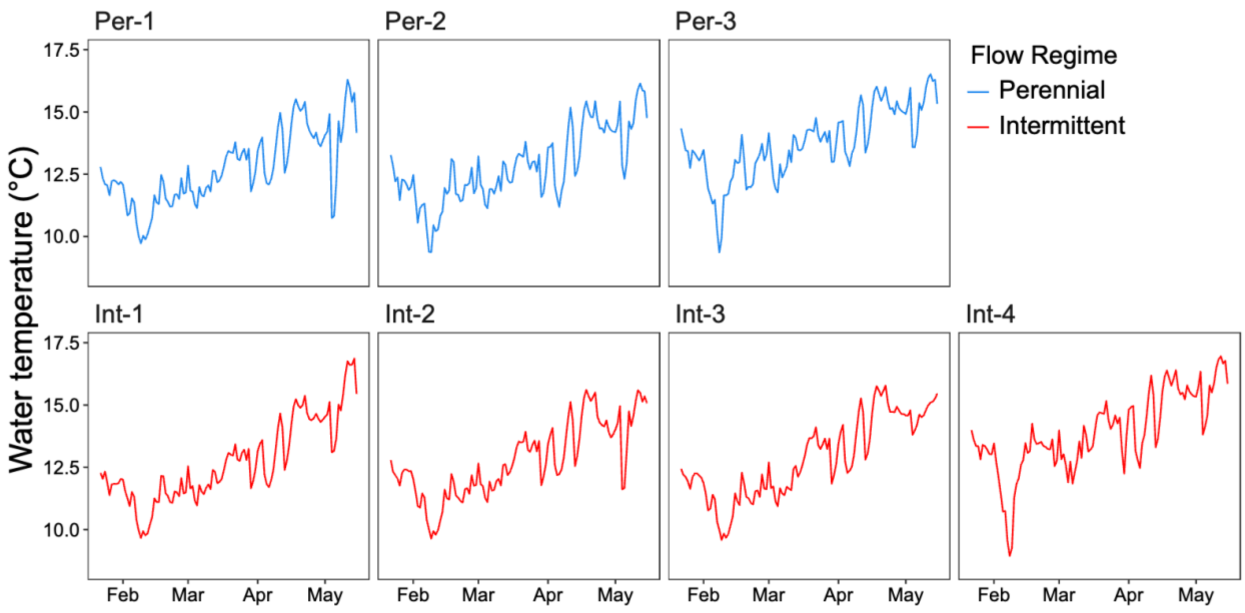


**Figure S4. Daily mean water temperature (°C) recorded at the seven study sites** during the decomposition experiment (January–May 2024). Perennial sites (blue, top row) and intermittent sites (red, bottom row) are ordered by site label. Temperature was recorded using dissolved oxygen and temperature loggers deployed at the upstream and downstream ends of each reach. Records from both sensors were pooled for sites with overlapping data (n = 5 sites), and data from a single sensor were used for the remaining two sites.

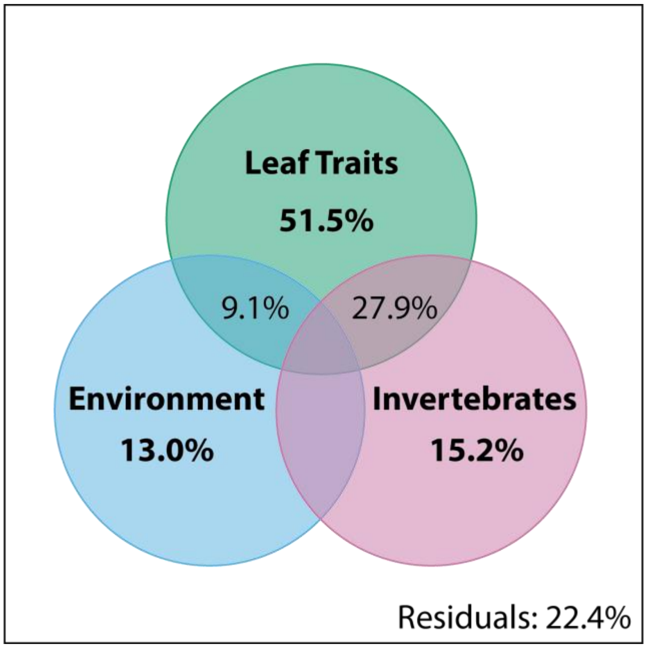


**Figure S5.** Variance partitioning of the total leaf litter decomposition rates among three predictor sets: litter quality ("Leaf Traits": specific leaf area and leaf nitrogen, phosphorus, and carbon content), invertebrate community metrics ("Invertebrates": densities of shredders, scrapers/grazers, predators, collectors, and parasites) and environmental variables ("Environment": flow regime, tributary, average water temperature, and drainage area). Values represent the percentage of variation (adjusted $R^2$) uniquely explained by each factor or shared between factors. Shared variance fractions <1% are not displayed.

**Table S1.** Characteristics of the seven study sites in Chalone Creek watershed, Pinnacles National Park, CA. Sites are ordered by flow permanence. Flow permanence represents the percentage of days with surface flow estimated from continuous Stream Temperature, Intermittency, and Conductivity (STIC) logger records from monitoring start date. Mean temperature reflects average daytime water temperature recorded during the decomposition experiment (January–May 2024).

| Site | Tributary | Flow Regime | Flow Permanence (%) | Monitoring Start | Drainage Area ($km^2$) | Mean Water Temperature (ºC) | Tree Species |
|---|---|---|---|---|---|---|---|
| Per-1 | Chalone Creek | Perennial | 100 | 2015 | 95.3 | 12.7 | PF, QA, SL |
| Per-2 | Sandy Creek | Perennial | 100 | 2015 | 57.3 | 12.3 | PF, QA, SL |
| Per-3 | Sandy Creek | Perennial | 100 | 2023 | 50.5 | 13.2 | PF, SL |
| Int-1 | Chalone Creek | Intermittent | 45.0 | 2015 | 116.8 | 12.7 | PF, QA, SL |
| Int-2 | Chalone Creek | Intermittent | 75.4 | 2023 | 144.2 | 12.4 | PF, SL |
| Int-3 | Chalone Creek | Intermittent | 38.6 | 2015 | 140.2 | 12.1 | PF, SL |
| Int-4 | Sandy Creek | Intermittent | 36.6 | 2015 | 44.5 | 13.6 | QA, SL |

Tree species abbreviations: PF = *Populus fremontii* (cottonwood), QA = *Quercus agrifolia* (oak), SL = *Salix laevigata* (willow).

**Table S2.** Invertebrate taxa identified from coarse mesh litter bags across all sites and leaf species. Total abundance is the sum of all individuals across all bags and sites ($n$ = 138). Frequency is the percentage of bags in which the taxon was present. Feeding group assignments follow Vieira et al. (2006); 1 = taxon assigned to feeding group, 0 = not assigned. Taxa marked with † are likely of terrestrial origin and were excluded from feeding group analyses.

| Order / Class | Family | Lowest Taxonomic Level | Total abundance | Frequency | Shredder | Collector / | Collector / | Scraper / | Predator | Parasite |
|---|---|---|---|---|---|---|---|---|---|---|
| Acari | | | 73 | 25% | 0 | 0 | 0 | 0 | 1 | 0 |
| Veneroida | Sphaeriidae | Pisidium | 1 | 1% | 0 | 0 | 1 | 0 | 0 | 0 |
| Hirudinea | Hirudinea | Hirudinea | 10 | 7% | 0 | 0 | 0 | 0 | 1 | 1 |
| Oligochaeta | | | 2,079 | 88% | 0 | 1 | 0 | 1 | 0 | 0 |
| Basommatophora | Physidae | Physa | 191 | 26% | 0 | 0 | 0 | 1 | 0 | 0 |
| Basommatophora | Planorbidae | Gyraulus | 7 | 5% | 0 | 0 | 0 | 1 | 0 | 0 |
| Copepoda | | | 16 | 7% | 0 | 0 | 1 | 0 | 0 | 0 |
| Coleoptera | Dytiscidae | | 36 | 15% | 0 | 0 | 0 | 0 | 1 | 0 |
| Coleoptera | Elateroidea | | 1 | 1% | 0 | 0 | 0 | 0 | 1 | 0 |
| Coleoptera | Elmidae | | 8 | 3% | 1 | 1 | 0 | 1 | 0 | 0 |
| Coleoptera | Haliplidae | | 9 | 2% | 1 | 0 | 0 | 1 | 0 | 0 |
| Coleoptera | Hydrophilidae | Hydrophilidae | 33 | 10% | 0 | 0 | 0 | 1 | 1 | 0 |
| Coleoptera | Hydrophilidae | | 9 | 4% | 0 | 0 | 0 | 1 | 1 | 0 |
| Coleoptera | Scirtidae | | 45 | 3% | 1 | 1 | 0 | 1 | 0 | 0 |
| Diptera | Ceratopogonidae | Bezzia | 172 | 43% | 0 | 0 | 0 | 0 | 1 | 0 |
| Diptera | Ceratopogonidae | Ceratopogon | 73 | 21% | 0 | 0 | 0 | 0 | 1 | 0 |
| Diptera | Ceratopogonidae | | 5 | 1% | 0 | 0 | 0 | 0 | 1 | 0 |
| Diptera | Chironomidae | Chironomidae (pupae) | 92 | 32% | 0 | 1 | 1 | 0 | 1 | 0 |

| | | | | | | | | | | |
|---|---|---|---|---|---|---|---|---|---|---|
| Diptera | Chironomidae | Chironominae | 5 | 2% | 0 | 1 | 1 | 0 | 0 | 0 |
| Diptera | Chironomidae | Chironomini | 713 | 60% | 0 | 1 | 0 | 0 | 0 | 0 |
| Diptera | Chironomidae | Diamesinae | 3 | 1% | 0 | 1 | 0 | 0 | 0 | 0 |
| Diptera | Chironomidae | Orthocladiinae | 1,315 | 66% | 0 | 1 | 0 | 1 | 0 | 0 |
| Diptera | Chironomidae | Podonominae | 5 | 2% | 0 | 1 | 0 | 0 | 0 | 0 |
| Diptera | Chironomidae | Tanypodinae | 899 | 70% | 0 | 0 | 0 | 0 | 1 | 0 |
| Diptera | Chironomidae | Tanytarsini | 3,754 | 52% | 0 | 1 | 1 | 0 | 0 | 0 |
| Diptera | Chironomidae | | 149 | 7% | 0 | 1 | 1 | 0 | 1 | 0 |
| Diptera | Dolichopodidae | | 4 | 2% | 0 | 0 | 0 | 0 | 1 | 0 |
| Diptera | Empididae | Clinocera | 5 | 3% | 0 | 0 | 0 | 0 | 1 | 0 |
| Diptera | Empididae | Empididae (pupae) | 1 | 1% | 0 | 0 | 0 | 0 | 1 | 0 |
| Diptera | Empididae | Trichoclinocera | 20 | 3% | 0 | 0 | 0 | 0 | 1 | 0 |
| Diptera | Ephydridae | | 3 | 1% | 0 | 0 | 0 | 0 | 1 | 0 |
| Diptera | Limoniidae | Limnophila | 3 | 1% | 0 | 0 | 0 | 0 | 1 | 0 |
| Diptera | Limoniidae | Limoniinae | 4 | 3% | 1 | 1 | 0 | 0 | 1 | 0 |
| Diptera | Limoniidae | Ormosia | 3 | 2% | 0 | 1 | 0 | 0 | 0 | 0 |
| Diptera | Muscidae | | 3 | 1% | 0 | 0 | 0 | 0 | 1 | 0 |
| Diptera | Pediciidae | Dicranota | 21 | 9% | 0 | 0 | 0 | 0 | 1 | 0 |
| Diptera | Psychodidae | | 5 | 1% | 0 | 1 | 0 | 0 | 0 | 0 |
| Diptera | Simuliidae | Simuliidae (pupae) | 17 | 7% | 0 | 0 | 1 | 0 | 0 | 0 |
| Diptera | Simuliidae | | 120 | 19% | 0 | 0 | 1 | 0 | 0 | 0 |
| Diptera | Stratiomyidae | Caloparyphus | 4 | 2% | 0 | 1 | 0 | 0 | 0 | 0 |
| Diptera | Stratiomyidae | Oxycera | 1 | 1% | 0 | 0 | 0 | 1 | 0 | 0 |
| Diptera | Stratiomyidae | | 2 | 1% | 0 | 1 | 0 | 1 | 0 | 0 |
| Diptera | Tabanidae | Tabanus | 12 | 9% | 0 | 0 | 0 | 0 | 1 | 0 |
| Diptera | Tipulidae | Tipula | 40 | 17% | 1 | 0 | 0 | 0 | 0 | 0 |
| Ephemeroptera | Ameletidae | Ameletus | 7 | 4% | 0 | 1 | 0 | 1 | 0 | 0 |

| | | | | | | | | | | |
|---|---|---|---|---|---|---|---|---|---|---|
| Ephemeroptera | Baetidae | | 148 | 15% | 0 | 1 | 0 | 1 | 0 | 0 |
| Ephemeroptera | Heptageniidae | | 1 | 1% | 0 | 1 | 0 | 1 | 0 | 0 |
| Ephemeroptera | Leptohyphidae | Tricorythodes | 17 | 6% | 0 | 1 | 0 | 0 | 0 | 0 |
| Ephemeroptera | Leptophlebiidae | Paraleptophlebia | 188 | 21% | 0 | 1 | 0 | 0 | 0 | 0 |
| Hemiptera | Aphidoidea † | | 2 | 1% | 0 | 0 | 0 | 0 | 0 | 0 |
| Hymenoptera | Formicidae † | | 6 | 3% | 0 | 0 | 0 | 0 | 0 | 0 |
| Lepidoptera | Pyralidae | | 1 | 1% | 1 | 0 | 0 | 0 | 0 | 0 |
| Megaloptera | Corydalidae | Neohermes | 3 | 1% | 0 | 0 | 0 | 0 | 1 | 0 |
| Megaloptera | Sialidae | Sialis | 8 | 2% | 0 | 0 | 0 | 0 | 1 | 0 |
| Odonata | Coenagrionidae | Argia | 25 | 12% | 0 | 0 | 0 | 0 | 1 | 0 |
| Odonata | Coenagrionidae | | 1 | 1% | 0 | 0 | 0 | 0 | 1 | 0 |
| Plecoptera | Capniidae | | 40 | 10% | 1 | 0 | 0 | 0 | 0 | 0 |
| Plecoptera | Nemouridae | Malenka | 719 | 24% | 1 | 0 | 0 | 0 | 0 | 0 |
| Plecoptera | Nemouridae | Nemoura | 13 | 7% | 1 | 0 | 0 | 0 | 0 | 0 |
| Plecoptera | Perlodidae | Baumannella | 3 | 1% | 0 | 0 | 0 | 0 | 1 | 0 |
| Plecoptera | Taeniopterygidae | Taenionema | 4 | 2% | 1 | 0 | 0 | 0 | 0 | 0 |
| Plecoptera | | Unidentified (early instar) | 565 | 33% | 1 | 0 | 0 | 0 | 0 | 0 |
| Thysanoptera † | | | 2 | 1% | 0 | 0 | 0 | 0 | 0 | 0 |
| Trichoptera | Brachycentridae | Micrasema | 31 | 12% | 1 | 0 | 1 | 0 | 0 | 0 |
| Trichoptera | Hydropsychidae | Hydropsyche | 154 | 15% | 0 | 0 | 1 | 0 | 0 | 0 |
| Trichoptera | Hydroptilidae | Hydroptila | 13 | 4% | 0 | 0 | 0 | 1 | 0 | 0 |
| Trichoptera | Hydroptilidae | Ochrotrichia | 10 | 5% | 0 | 0 | 0 | 1 | 0 | 0 |
| Trichoptera | Lepidostomatidae | Lepidostoma | 41 | 14% | 1 | 0 | 0 | 0 | 0 | 0 |
| Trichoptera | Leptoceridae | Nectopsyche | 1 | 1% | 1 | 1 | 0 | 0 | 0 | 0 |
| Trichoptera | Limnephilidae | Limnephilus | 1 | 1% | 1 | 0 | 0 | 0 | 0 | 0 |
| Trichoptera | Philopotamidae | Wormaldia | 1 | 1% | 0 | 0 | 1 | 0 | 0 | 0 |

| | | | | | | | | | | |
|---|---|---|---|---|---|---|---|---|---|---|
| Trichoptera | Sericostomatidae | Gumaga | 6 | 2% | 1 | 1 | 0 | 1 | 0 | 0 |
| Insecta | | Unidentified (terrestrial) † | 31 | 14% | 0 | 0 | 0 | 0 | 0 | 0 |
| Amphipoda | Gammaridae | Gammarus | 29 | 7% | 1 | 1 | 0 | 1 | 0 | 0 |
| Amphipoda | Hyalellidae | Hyalella | 558 | 36% | 1 | 1 | 0 | 1 | 0 | 0 |
| Amphipoda | | | 29 | 12% | 1 | 1 | 0 | 1 | 0 | 0 |
| Isopoda | Armadillidae † | | 3 | 1% | 0 | 0 | 0 | 0 | 0 | 0 |
| Ostracoda | | | 88 | 28% | 0 | 0 | 1 | 0 | 0 | 0 |
| Turbellaria | | | 5 | 2% | 0 | 0 | 0 | 0 | 1 | 0 |

**Table S3.** Results of generalized linear mixed-effects models (GLMMs) testing effects of flow permanence and leaf species on invertebrate functional feeding group density and richness. All models used a negative binomial distribution with site as a random intercept. Flow permanence (%) was z-scored prior to model fitting. Bold values indicate $p < 0.05$.

| Response | | Flow permanence | | Leaf Species | | Flow × Leaf Species | |
|---|---|---|---|---|---|---|---|
| | | $\chi^2(1)$ | p-value | $\chi^2(2)$ | p-value | $\chi^2(2)$ | p-value |
| Shredder | Density | **5** | **0.025** | 3.15 | 0.207 | 0.245 | 0.885 |
| | Richness | 2.39 | 0.122 | **7.76** | **0.021** | 0.027 | 0.987 |
| Collector | Density | 1.6 | 0.206 | **7.42** | **0.024** | 0.631 | 0.729 |
| | Richness | 0 | 0.991 | 2.27 | 0.322 | 1.564 | 0.458 |
| Scraper | Density | 1.39 | 0.238 | *5.25* | 0.072 | 0.106 | 0.948 |
| | Richness | 0.002 | 0.962 | *5.58* | 0.062 | 0.42 | 0.811 |
| Predator | Density | 0.004 | 0.949 | **9.04** | **0.011** | 1.136 | 0.567 |
| | Richness | 0.12 | 0.727 | 3.35 | 0.188 | 0.383 | 0.826 |
